# Development of immortalized *Callithrix jacchus* kidney cell lines supporting infection with a panel of viruses

**DOI:** 10.1101/2025.11.13.688326

**Authors:** Sabine Gärtner, Pamela Stomberg, Stefan Pöhlmann, Michael Winkler

## Abstract

The common marmosets (Callithrix jacchus) are valuable non-human primate (NHP) animal models in biomedical research, including infectious diseases modelling. However, for in vitro studies only few immortalized cell lines have been generated, and additional lines are needed and will help to comply with the 3R principles or replacement, reduction and refinement. Here, we present the generation and characterization of three cell lines derived from kidney tissue, which were immortalized by transduction of SV40 large T antigen. The cell lines display an epithelioid morphology, show differential podoplanin expression and are likely of pericyte origin, as deduced from expression profiles of marker genes obtained by RNA sequencing analysis (RNA-seq). All cell lines had a functional interferon system, as shown by responsiveness to human IFNβ and marmoset IFNα14 and the induction of interferon-stimulated genes. Infection with retroviral pseudotypes demonstrated susceptibility to entry driven by glycoproteins from a wide range of human pathogenic viruses. Finally, these cell lines are highly permissive for Zika virus, for which marmosets are a model organism, and Herpes simplex virus 1, which causes a deadly disease in marmosets. We believe that these cell lines are a valuable resource for in vitro studies on marmosets.

## Introduction

The common marmosets (*Callithrix jacchus*) are small, neotropical, non-human primates (NHP) that have emerged as a valuable model organism in biomedical research, particularly in the fields of immunology, neuroscience, and infectious disease modeling [1–3]. Their small size, short lifespan, and close phylogenetic relationship to humans make marmosets an attractive alternative to larger primates, such as rhesus macaques, as well as to traditional rodent models like mice and rats. Unlike rhesus macaques, marmosets do not carry pathogens harmful to humans, such as Macacine alphaherpesvirus 1 (Herpes B virus) [3]. Conversely, marmosets are naturally susceptible to numerous human pathogenic viruses [1, 3, 4]. This includes highly pathogenic viruses, such as Lassa virus (LASV) [5, 6], Ebola virus (EBOV) and Marburg virus (MARV) [7], emerging viruses, such as Zika virus (ZIKV) [8, 9], or the respiratory pathogens SARS-CoV-1 [10], SARS-CoV-2 [11] and MERS-CoV [12], but also herpesviruses such as Epstein-Barr virus [13] and Herpes simplex virus 1 (HSV-1), which can cause life threatening infections in common marmosets [14–16].

The use of animals in research is strictly regulated by laws, and animal experiments need to be conducted under an ethical framework, which includes adherence to the 3R principles of replacement, reduction and refinement [17]. Immortalized, well-characterized cell lines are powerful tools that can be integrated into experimental workflows, for example to evaluate virus entry or innate immune responses in vitro, and thus help to comply with the 3R principles. However, for marmosets, in contrast to humans, only a small number of cell lines have been described, including immortalized fibroblasts [18, 19], ovarian granulosa and theca cells [20] and hepatic progenitor cells [21], with a single marmoset cell line being available from ATCC. Therefore, more immortalized cell lines are needed, particularly for in vitro studies in virology, oncology and drug metabolism [22].

In this study, we describe the establishment and characterization of three immortalized cell lines from marmoset kidney tissue, which will serve as a valuable resource for the marmoset research community. The cell lines are highly related to pericytes as evidenced by RNAseq analysis and retained a functional interferon (IFN) system. The cell lines were susceptible to the entry of pseudoparticles bearing the glycoproteins of multiple highly pathogenic viruses and supported growth of the RNA virus ZIKV as well as the DNA virus HSV-1.

## Materials and Methods

### Plasmids and oligonucleotides

The previously described plasmids HIV gag-pol(SCA), pLenti-IP-large T, MLV-gag/pol and MLV-luc were used for lentiviral or retroviral transduction [23–25]. Additionally, expression plasmids for various viral glycoproteins were used, including those from vesicular stomatitis virus (*Vesiculovirus indiana,* VSV-G), Ebola virus (*Orthoebolavirus zairense,* EBOV-GP), Marburg virus (*Orthomarburgvirus marburgense,* MARV-GP), influenza A virus strain WSN (*Alphainfluenzavirus influenza*, HA and NA), Lassa virus (*Mammarenavirus lassaense*, LASV-GP), Machupo virus (*Mammarenavirus machupoense*, MACV-GP), Lymphocytic choriomeningitis virus strain Armstrong (*Mammarenavirus choriomeningitidis,* LCMV-GP), La Crosse virus (*Orthobunyavirus lacrosseense,* LACV-GPC), Rift Valley fever virus (*Phlebovirus riftense,* RVFV-Gn/Gc) or severe fever with thrombocytopenia syndrome virus (*Bandavirus dabieense,* SFTSV-Gn/Gc) have been previously described [26–33].

Expression plasmids for IFNs were newly generated. For this, human IFNB1 was amplified from genomic DNA of A549 cells using primer hIFNb1-5Acc (5-CCGGTACCATGACCAACAAGTGTCTCC-3)/ hIFNb1-3Bgl (5-GGAGATCTTCAGTTTCGGAGG-3) and cloned into pCAGGS [34] via Acc65I and BglII to obtain pCAGGS-hIFNB1. *Callithrix* IFNa14 was amplified from genomic DNA of CajaK-D4 cells using primer cjIFNa2-5Acc (5-CCGGTACCATGRCCTTGACCTTTCCTTTA-3)/cjIFNa2-3Sal (5-CCCGTCGACTCATTTCTTACTTCTTAAMCC-3) and cloned into pCAGGS via Acc65I and SalI to obtain pCAGGS-cjIFNA14. This clone has an amino acid exchange A161T with respect to the cjIFNa14 sequence (LOC100406466; XM_002763506.4).

### Cell culture

Cell lines 293T (human kidney, female; DSMZ catalogue no. ACC-635, RRID:CVCL 0063), Vero76 (African green monkey kidney, female; ATCC catalogue no. CRL-1586; RRID: CVCL 0574; kindly provided by Andrea Maisner) and A549 (human lung, male; CRM-CCL-185, ATCC, RRID:CVCL_0023; kindly provided by Benjamin tenOever) were cultured in DMEM supplemented with 10% fetal calf serum (FCS) and penicillin/streptomycin. Human cell line identity was verified through short tandem repeat (STR) analysis [35], and regular mycoplasma contamination screenings were performed on all cell lines.

### Establishment of primary cultures

Primary cultures were established from kidney tissue obtained from a 12-year and 10-month-old male animal. The tissue was finely chopped into small pieces (<1mm) and 2-3 pieces were transferred to Eppendorf tubes containing a 50 µl digestion mixture with 5 mg collagenase IV. After incubation on a shaker (37°C, 800rpm) for 1-1.5 h, cell suspensions were transferred into 6-well plates containing DMEM supplemented with 10% FCS and antibiotics (penicillin, streptomycin, gentamycin, nystatin and amphotericin B).

### Retroviral transduction

For immortalization, cells were transduced with a lentiviral vector expressing SV40 large T antigen. For preparation of transducing particles, 293T cells were seeded in T25 flasks and transfected with a mixture of 6 μg lentiviral vector pLenti-IP-large T, 3 μg HIV gag-pol(SCA) and 3 μg pHIT/G expressing VSV-G, using the calcium phosphate method. Transduced cells were subsequently selected with 1 µg/mL puromycin. Established cell lines were named as ***Ca****llithrix **ja**cchus* **k**idney (CajaK) cells with a suffix for distinction (CajaK-C3, CajaK-D4, CajaK-E5).

### Microscopy

Brightfield images were taken of cells grown in 24-well plates using a ZOE cell imaging system (Bio-Rad, Feldkirchen, Germany) at 20x magnification. Images were processed (cropping, scale bar) using Fiji/ImageJ 1.54f [36].

### RNAseq analysis

Cells were seeded in 6-well plates at a density of 250,000 cells/well. After overnight growth, total cellular RNA was isolated using Trizol reagent (Ambion, Carlsbad, CA, USA) following the manufacturer’s instructions. The isolated RNA was subjected to 50 bp single-end bulk RNA sequencing (Illumina HiSeq 4000) at the Core Unit NGS-Integrative Genomics (NIG), University Medical Center Göttingen (UMG). The NIG provided raw count profiles and normalized TPM values. To visualize the expression of marker genes, heatmap graphs were generated using the R platform (version 4.5.1; https://www.R-project.org/) and the pheatmap package [37, 38]. Marker genes were chosen based on a published single-cell RNA sequencing (scRNAseq) analysis of human adult kidney samples [39]. For our analyses, we used canonical marker genes (Suppl. Fig. 3B in [39]) and a panel of genes specifically expressed in pericyte-2 cells or fibroblasts (Suppl. Table 3 in [39]). In the latter case, only genes with highest significance and expression in our cell line samples were selected.

### Flow cytometry

The CajaK cell lines generated here were seeded in 6-well plates at 200,000 cells/well and incubated for two days. The cells were then detached using PBS containing 5 mM EDTA. After centrifugation (1,200 rpm, 5 min) cell pellets were washed in FCS buffer (PBS, 5% FCS, 2 mM EDTA) and reacted with either rat anti-podoplanin antibody (1:100; Origene AM01133PU-N, Rockville, MD, USA) or rat IgG2a isotype control antibody (Antikoerper-online ABIN5675832, Aachen, Germany) in a volume of 50 µL for 30 min on ice. After three washes in FCS buffer, cells were incubated (30 min, room temperature) with Alexafluor488-conjugated donkey anti-rat antibody (1:100; Invitrogen A21208, Carlsbad, CA, USA), followed by three final washes in FCS buffer and fixation in 2% PFA. Samples were analyzed on an ID7000 Spectral Cell Analyzer (Sony, San Jose, CA, USA). Diagrams for the figures were prepared using EasyFlowQ version 1.7 [40].

### Quantitative real-time PCR

CajaK cell lines and A549 control cells were seeded in 12-well plates at 150,000 cells/well. The next day, the cells were either left untreated, treated with pan-IFN (PBL Assay Science 11200-2, Piscataway, NJ, USA) at 100 U/mL, or infected with VSV ncp* at MOI 0.1. After 24 h cells were harvested in Trizol (Ambion, Carlsbad, CA, USA) and RNA purified according to instructions of the manufacturer. For each sample, 1 µg RNA was treated with 0.5 U RNase-free DNase I for 10 min at 37°C and stopped with EDTA. After denaturation at 75°C (10 min), cDNA synthesis was performed with random hexamers using the SuperScript™ III First-Strand Synthesis System (Thermo, Waltham, MA, USA). For quantitative PCR analysis, 1µL of cDNA and primers against IFNB1 (forward 5-CAGCAATTTTCAGTGTCAGAAGC-3, reverse 5-TCATCCTGTCCTTGAGGCAGT-3), MX1 (forward 5-TTCAGCACCTGATGGCCTATC-3, reverse 5-TGGATGATCAAAGGGATGTGG-3), and the housekeeping gene 18S rRNA (forward 5-GATCCATTGGAGGGCAAGTCT-3, reverse 5-CCAAGATCCAACTACGAGCTT-3) were used [41–43]. The reactions were performed on a Rotorgene Q platform (Qiagen, Hilden, Germany) using the QuantiTect SYBR Green PCR kit (Qiagen, Hilden, Germany). Calculation of IFNB1 and MX1 induction was done using the 2^-ΔΔCT^ method using 18S rRNA as reference [44].

### Conditioned IFN supernatants

For generation of conditioned supernatants, 293T cells were seeded in T25 flaks at 1,500,000 cells/flask. The next day, cells were transfected with 5 µg of pCAGGS-hIFNB1 or pCAGGS-cjIFNa14 using polyethyleneimine (PEI) [45]. For transfection, 12 µg DNA and 36 µL of PEI (Polyscience, Inc., 23966; 0.1 % w/v) were separately diluted in 250 µL OptiMEM, mixed and incubated at room temperature before adding the mixtures to the cells. After overnight incubation, medium was replaced with fresh medium (DMEM containing 10% FCS) and cells were incubated for additional 48 h. Cell culture supernatant was collected, filtered through 0.2 µm filters, aliquoted and stored at −80°C.

Interferon activity was determined using a VSV strain essentially as described in Berger Rentsch and Zimmer [46]. Briefly, both A549 and CajaK-E5 cells were seeded in 96-well plates at 10,000 cells/well and incubated overnight. Two-fold dilution series were prepared for IFN-containing supernatants as well as for pan-IFN (starting concentration 400 U/mL) and cells treated with 100 µL/well for 24 h. On the next day, medium was removed and cells were infected with VSV*ΔG(Fluc) at MOI 5 (100 µL volume). After 6 h incubation cells were lysed in 50 μl Luciferase Cell Culture Lysis Reagent (Promega, Madison, WI, USA) for 20 min at room temperature. The lysates were transferred to white 96-well plates, mixed with 50 µL Beetle Juice reagent (PJK, Kleinblittersdorf, Germany) and luciferase activity measured in a Hidex Sense luminescence reader (Hidex, Turku, Finland). After subtraction of background (untreated, uninfected cells) the infection rate was calculated. Assuming 1 unit IFN resulting in 50% inhibition, the activities in the sample solutions (titrated on CajaK-E5 cells) were calculated. Titration of pan-IFN on A549 cells served as control.

### Immunoblot

For analysis of large T expression after IFN treatment, CajaK cell lines, A549 and 293T control cells were seeded in 6-well plates at 250,000 cells per well. After 48 h, the cells were lysed in SDS sample buffer (30 mM Tris pH 6.8, 1 mM EDTA, 10% glycerol, 2% SDS, 0.1% bromophenol blue, 5% beta-mercaptoethanol), heated to 95°C for 15 min and separated using SDS-PAGE. The proteins were transferred onto nitrocellulose membranes (Protran; Amersham, Freiburg, Germany) using a Mini-Protean electrophoresis system (Bio-Rad, Hercules, CA, USA). Filters were blocked in PBS-T/MP (PBS with 0.1% Tween-20 and 5% skim milk powder) for 1 h at room temperature followed by incubation with primary antibodies overnight at 4°C. Primary antibodies mouse anti-large T monoclonal PAb108 (BD Biosciences 554150, San Diego, CA, USA) and mouse anti-beta-actin (Sigma A5441, Taufkirchen, Germany) were diluted 1:1,000 in PBS-T/MP, respectively. After washing three times in PBS-T (PBS with 0.1% Tween-20), filters were incubated for 1 h at room temperature with horseradish conjugated anti-mouse antibodies (Dianova, Hamburg, Germany) diluted 1:10,000 in PBS-T/MP. After three final washes in PBS-T, filters were incubated with Westar Antares chemiluminescence substrate (Cyanagen XLS0142,0250; Bologna, Italy) and imaged using a Azure 600 Imager (Azure Biosystems, Singapore).

For analysis of IFITM3 expression after IFN treatment, CajaK cell lines and A549 control cells were seeded in 24-well plates at 50,000 cells per well. The next day, the cells were treated with conditioned medium of either hIFNB1 or cjIFNa14 at a concentration of 1,000 U/mL for 24 h. After treatment, the cells were lysed in SDS sample buffer heated to 95°C for 5 min and separated using SDS-PAGE (15%). After transfer and blocking, filters were processed as described above, using primary antibodies rabbit anti-IFITM (Abgent AP1153a, San Diego, CA, USA) and mouse anti-beta-actin (Sigma A5441, Taufkirchen, Germany) diluted 1:1,000 and 1:500 and horseradish conjugated secondary antibodies anti-mouse or anti-rabbit (Dianova, Hamburg, Germany) diluted 1:10,000. For detection, filters were incubated with HRP Juice (PJK Biotech, Kleinblittersdorf, Germany) and imaged using a ChemoCam Imager (Intas, Göttingen, Germany).

### Transduction with retroviral pseudoparticles

Production of retroviral pseudoparticles bearing glycoproteins of multiple viruses followed established protocols [24, 25, 47, 48]. Briefly, 293T cells were seeded into T25 flasks and transfected with 6 μg vector MLV-luc, 3 μg MLV-gag-pol and 3 μg expression plasmid for the respective viral glycoprotein. After two days, cell culture supernatants were harvested, filtered through 0.45 µm filters, aliquoted and stored at −80°C. Transduction efficiency was first quantified on 293T cells by measuring luciferase activity.

For comparative experiments, CajaK cell lines were seeded in 96-well plates at 10,000 cells/well in 50 µL medium. Before infection, pseudoparticle preparations were adjusted for comparable luciferase yield as measured in 293T cells. For infection, 50 µl adjusted pseudoparticle supernatant was added to the cells, incubated for 4-6 h until an additional 100 µL medium was added. After 3 days, cells were lysed using 50 μl Luciferase Cell Culture Lysis Reagent (Promega, Madison, WI, USA) for 20 min at room temperature. After transfer to white 96 well plates, samples were mixed with 50 µL Beetle Juice reagent (PJK, Kleinblittersdorf, Germany) and luciferase activity measured in a Hidex Sense luminescence reader (Hidex, Turku, Finland).

### Virus

Zika Virus (*Orthoflavivirus zikaense,* ZIKV) strain SPH2015 has been previously cloned from RNA and rescued from plasmid in Vero76 cells [49]. Herpes simplex virus 1 (*Simplexvirus humanalpha1,* HSV-1) strain F was rescued from bacterial artificial chromosome (BAC) pHSVF-CREin (kindly provided by Greg Smith) [50]. To obtain a HSV-1 reportervirus expressing Gaussia luciferase (HSV-1-cmvGluc), an expression cassette consisting of human cytomegalovirus enhancer-promoter, Gaussia luciferase and bovine growth hormone poly A signal present in pcDNA3-Gluc-en [51] was amplified using primers ep-HSV1-UL3BpA-5 (5-CACTGCCCGTCGCGCGTGTTTGATGTTAATAAATAACACATAAATTTGGCTCCCCAGCATGCCTGCTATT-3) and ep-HSV1-UL4cmv-3 (5-AAATGCCCCCCCCCCCTTGCGGGCGGTCCATTAAAGACAACAAACAACCAGTTGACATTGATTATTGACT-3). The purified PCR product was electroporated into E. coli GS1783 containing the BAC pHSVF-CREin and integrated using recombineering as previously described [50–52], followed by rescue on Vero76 cell. The Papiine alphaherpesvirus 2 (*Simplexvirus papiinealpha2,* PaHV-2) containing Gaussia luciferase reporter gene driven by the human cytomegalovirus enhancer-promoter (PaHV-2-cmvGluc) was previously described [51]. Likewise, the Cercopithecine alphaherpesvirus 2 (*Simplexvirus cercopithecinealpha2,* CeHV-2) containing Gaussia luciferase linked via a 2A sequence to ICP4 (CeHV-2-ICP4-Gluc) has been described [52].

### Virus replication kinetics and titration

For single step growth curves, CajaK cell lines or Vero76 control cells were seeded in 24-well plates at 50,000 cells/well and infected on the next day with either ZIKV or HSV-1 at MOI 1. For infection, medium was removed and inoculum added in triplicates, incubated for 1 h at 37°C and replaced with fresh medium. After 1, 24, 48 and 72 h, supernatants were harvested, centrifuged at 4,000 rpm for 5 min and the cleared supernatants frozen at −80°C. Titration of HSV-1 by plaque assay and ZIKV by focus forming assay, were performed on Vero76 as previously described [24, 52].

For multicycle growth experiments, marmoset cell lines were seeded in 6-well plates at 250,000 cells. On the next day, cells were infected with HSV-1-cmvGluc, PaHV-2-cmvGluc or CeHV-2-ICP4-Gluc in triplicates with 100 pfu/well (MOI 0.0004). At time intervals of 6-18 h aliquots of 25 µl were collected until 96 h postinfection and stored at –20°C. After collection of all samples, luciferase activity was measured in a Hidex sense plate reader (Hidex, Turku, Finland) using coelenterazine (PJK, Kleinblittersdorf, Germany) as a substrate, diluted to a final concentration of 1.5 mM in D-PBS (with Ca and Mg) [53].

### Statistical analysis

For statistical analyses and preparation of graphs GraphPad Prism software (version 10.6.1; GraphPad Software, Boston, MA, USA) was used. Significance was tested by one-way ANOVA with Dunnett’s correction for multiple comparisons assuming a lognormal distribution [54].

### Ethical statement

The German Primate Center is a non-profit independent research and service institute since 1977. It is registered and authorised by the local and regional veterinary governmental authorities (Ref. no.122910.3311900, PK 36674). All works regarding animals, animal reseach and sampling followed valid national and international guidelines. Good veterinary practice was applied to all procedures whenever animals are handled. According to the Animal Welfare Committee, the post-mortem use of samples from non-human primates derived from animals with a permit in accordance with §4, §7 and §11 of the Animal Welfare Act is in line with the German legal and ethical requirements for appropriate animal experiments with non-human primates. The consultation of the Animal Welfare Committee and the Animal Welfare Officer is documented under no. E3-25.

## Results

### Establishment of immortalized marmoset kidney cell line

To establish marmoset kidney cell lines, we obtained leftover material from a male animal aged 12 years and 10 months that had been sacrificed as part of an unrelated study (Fig. 1a). The kidney tissue sample was dissected into small fragments and cultivated in vitro until cultures of growing adherent cells were established. These cells were then transduced with a lentiviral vector expressing SV40 large T antigen and subjected to puromycin selection [25]. Individual puromycin-resistant colonies were picked and grown as individual cultures and named as ***Ca****llithrix **ja**cchus* **k**idney (CajaK) cells with a suffix for distinction (CajaK-C3, CajaK-D4, CajaK-E5). Morphological examination revealed that the CajaK cells exhibited an epithelioid morphology, with CajaK-D4 cells displaying a more spread-out appearance and CajaK-C3 and CajaK-E5 cells showing a more rounded morphology (Fig. 1b). All cell lines expressed the immortalization marker large T antigen, as demonstrated by immunoblot (Fig. 1c). Further analysis of these cell lines revealed differential expression of the kidney marker podoplanin (Fig. 1d), with CajaK-E5 cells exhibiting clear surface expression, CajaK-C3 cells showing no expression, and CajaK-D4 cells displaying an intermediate phenotype. This pattern was corroborated by subsequent RNAseq analysis (Fig. 1e).

**Figure 1:**
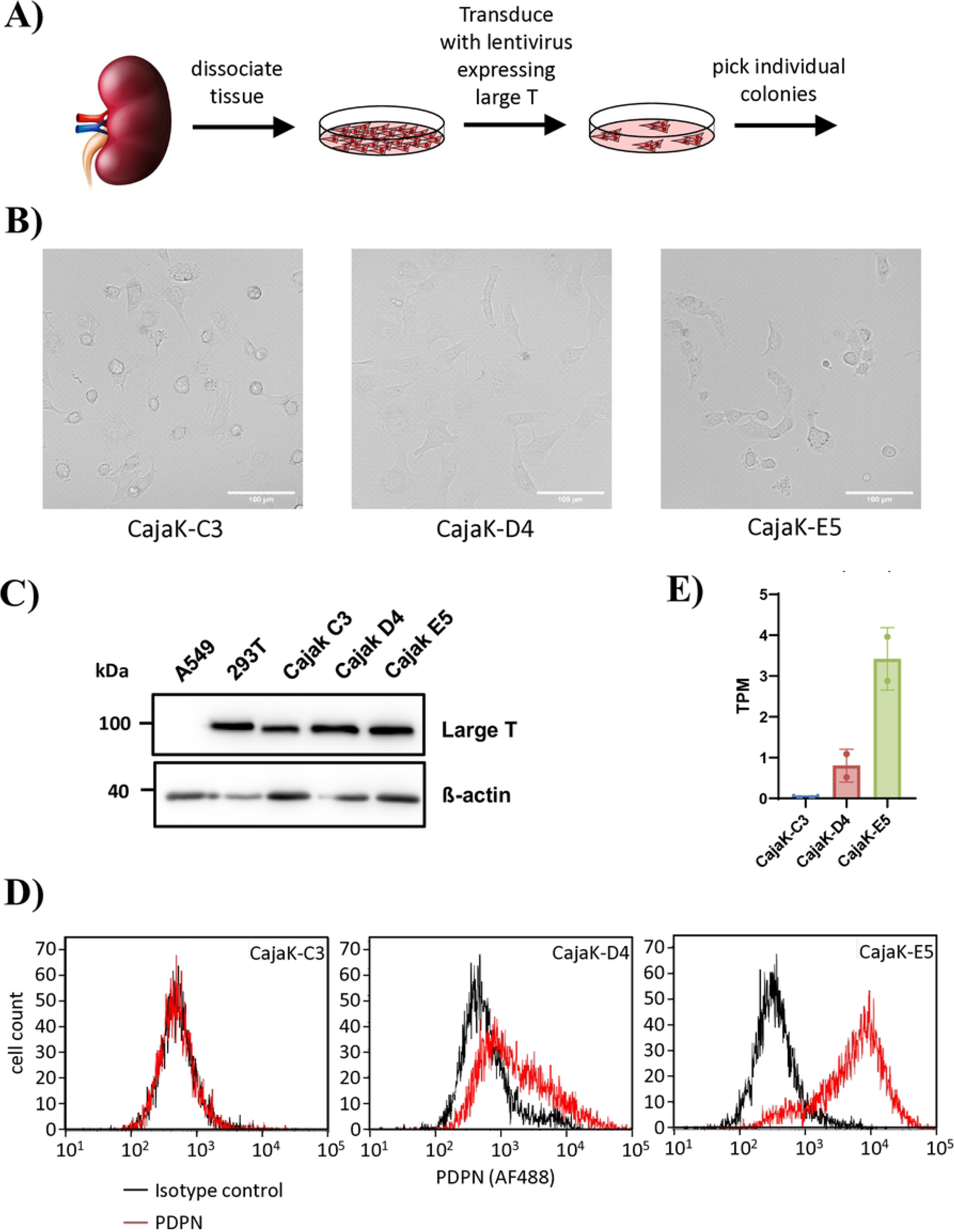
Generation and initial characterization of marmoset kidney cell lines. (A) Overview of the strategy used to generate marmoset kidney cell lines (image of kidney designed by brgfx/Freepik). (B) Morphology of marmoset kidney cell lines. Cell lines were seeded in 24-well plates and bright field images were taken at 20x magnification using a ZOE cell imaging system. Scale bars indicate 100 µm. (C) Immunoblot analysis of expression of immortalization marker large T in marmoset kidney cell lines. A549 and 293T cells served as negative or positive control, respectively. Beta-actin served as loading control. Results were confirmed in a separate experiment. (D) Analysis of podoplanin (PDPN) surface expression by flow cytometry. Cell lines were detached and stained with anti-podoplanin antibody (red) or isotype control antibody (black). Data were filtered to include only intact singlet cells. The results were verified through a separate, independent experiment. (E) Comparison of PDPN expression at transcriptional level. Cell lines were subjected to RNAseq, transcripts per million (TPM) are shown.

### Marmoset kidney cell lines display pericyte identity

For analysis of cell identity, total RNA was isolated from duplicate samples and subjected to RNA sequencing (RNAseq) (Suppl. Table 2). Principal component analysis demonstrated distinct identities for each cell line (Suppl. Fig. 1). To infer the cell type identity, we first analyzed expression of canonical marker genes based on scRNAseq analysis of human adult kidney samples [39]. As shown in Fig. 2a, all the cell lines demonstrated strong expression of the pericyte marker COX4I2 (Suppl. Table 3). Using panels of genes specifically expressed in individual cell types, we confirmed medium to high expression of genes specific to pericytes (Fig. 2b), while genes specific for fibroblasts typically showed low expression in our cell lines (Suppl. Tabls 4-5). Thus, RNAseq analysis indicates that our immortalized cell lines were likely derived from cells of pericyte origin.

**Figure 2:**
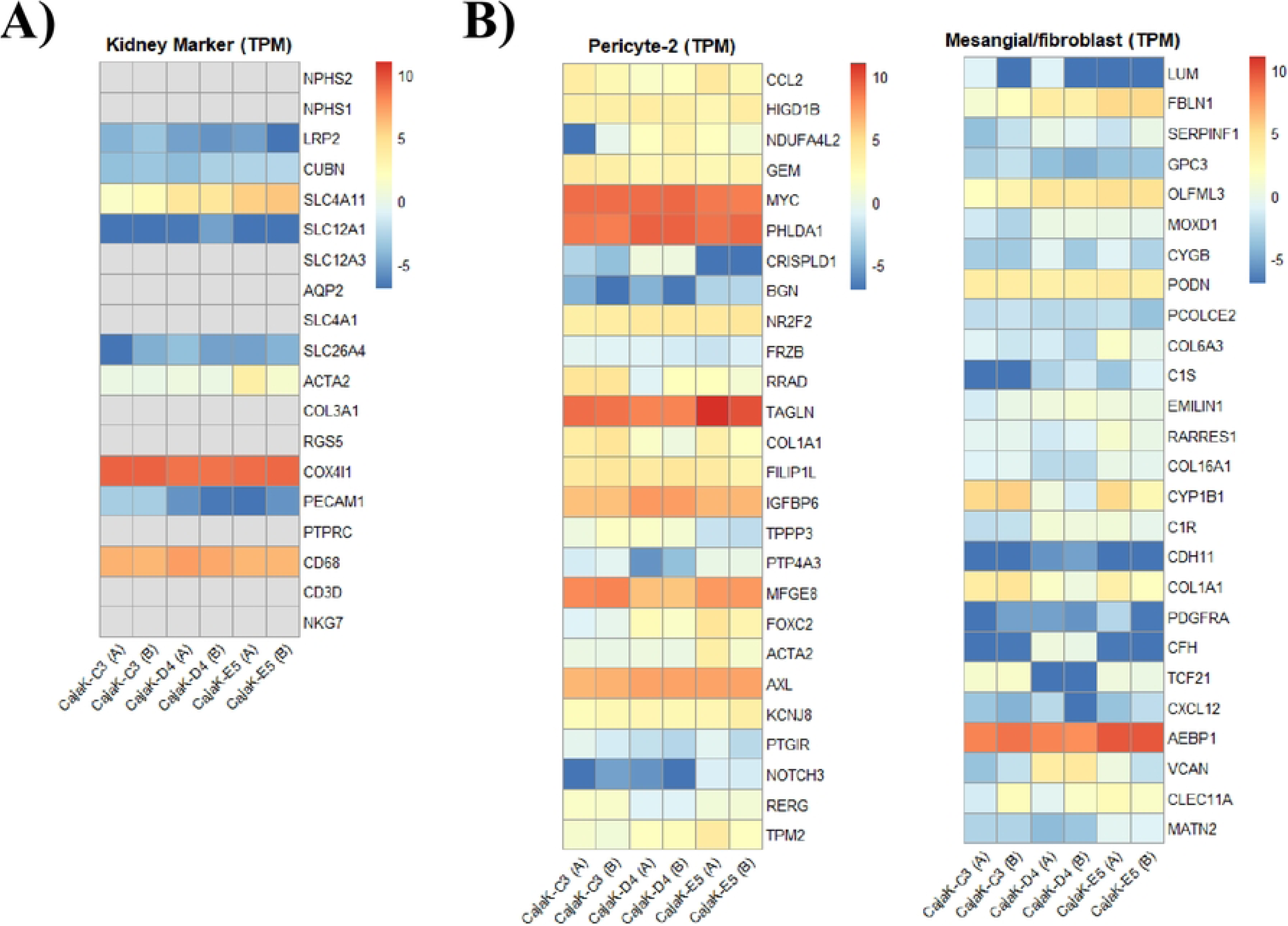
Cell type identification by transcriptome sequencing. Marmoset kidney cell lines were subjected to RNAseq analysis. For each cell line total RNA was isolated from two independent samples. (A) Expression in marmoset kidney cell lines of canonical markers for human kidney cell types as defined by Subramanian et al, 2019 Suppl. Fig. 3B [39]. Shown is a heatmap of log2-transformed TPM data derived from marmoset kidney cell lines. Grey boxes indicate that no signals were obtained. (B) Expression in marmoset kidney cell lines of genes selectively expressed in pericytes (left panel) and fibroblasts (right panel), as defined by Subramanian et al, 2019 Suppl. Table. 3 [39]. Shown are heatmaps of log2-transformed TPM data derived from marmoset kidney cell lines.

### Supplementary Figure 1: Principal component analysis

Principal component analysis based on RNA-seq profiles of cell lines CajaK-C3 (red), CajaK-D4 (green) and CajaK-E5 (blue). Duplicate samples are labelled A and B.

### Immortalized marmoset kidney cells lines have a functional IFN system

Next, we characterized whether the marmoset cell lines expressed interferon β (IFNB1) or the interferon stimulated gene (ISG) MX1 in response to treatment with IFN α or infection with vesicular stomatitis virus (VSV). The human lung adenocarcinoma cell line A549 has a functional IFN system and was used as positive control [24, 55]. Cells were incubated for 24 h with either pan-IFN, a chimeric human IFN α, or VSV-ncp*, a VSV variant which strongly induces IFN [56]. Induction of IFNB1 and MX1 was analyzed by quantitative RT-PCR. Infection with VSV-ncp* clearly induced both IFNB1 and MX1 (Fig. 3a), albeit MX1 induction was lower in the marmoset cell lines than in A549 cells. Treatment with pan-IFN did not induce IFNB1, as expected [24], but unexpectedly also did not induce MX1 in marmoset cell lines, while a strong induction was measured in A549 cells. This unresponsiveness of marmoset cells to human IFN α is in agreement with a previous report on marmoset hepatocytes [57].

**Figure 3:**
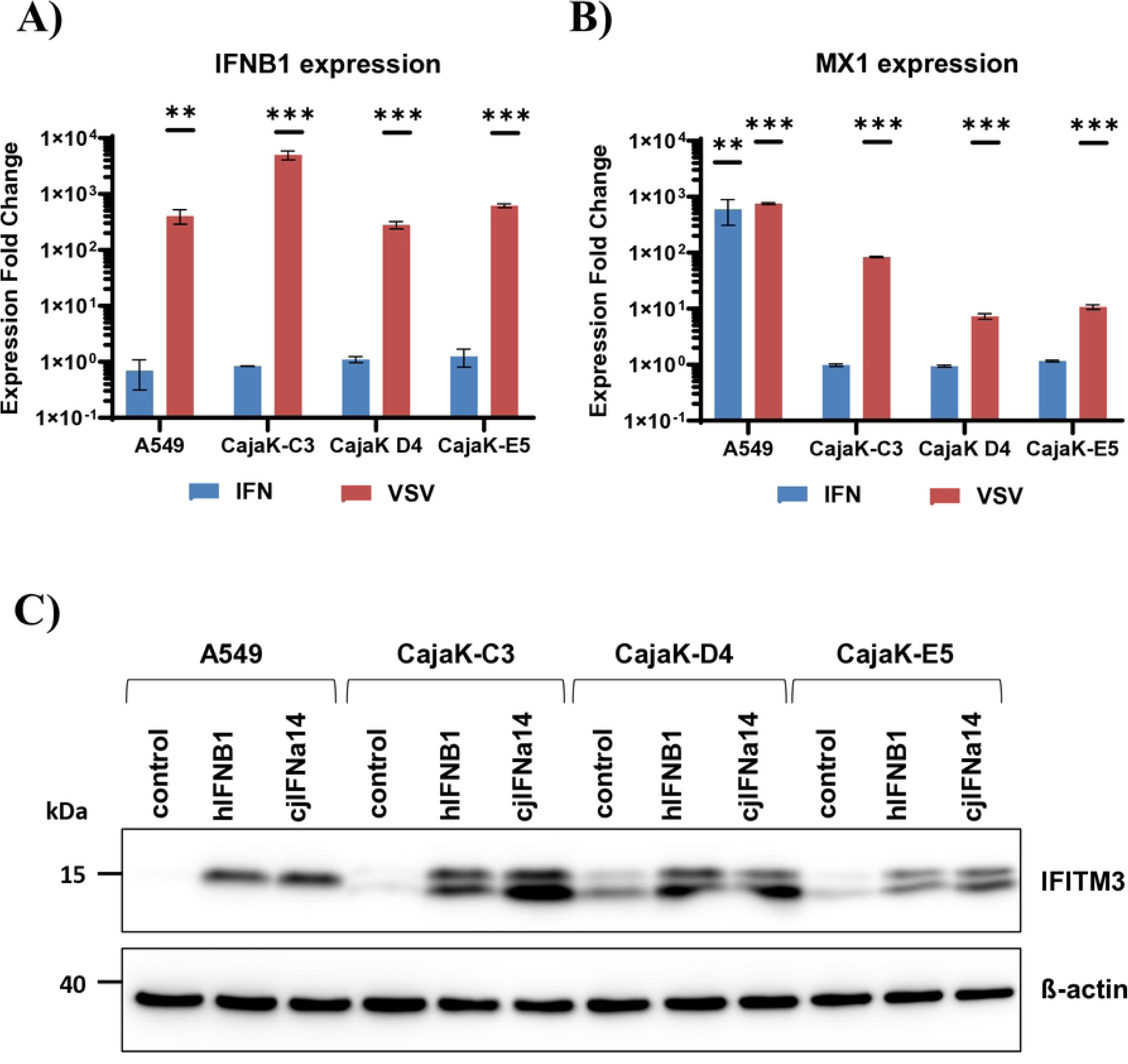
Immortalized marmoset kidney cells respond to virus infection and IFN. (A, B) Marmoset kidney cell lines or control cells (A549) were treated with pan-IFN (100 U/mL) or infected with vesicular stomatitis virus (VSV ncp *; MOI 0.1) for 24 h. Untreated cells served as control. Total RNA was isolated and expression of interferon beta (IFNB1) (A) or interferon-stimulated gene MX1 (B) quantified by qRT-PCR. Transcript levels were normalized against 18S rRNA and expression fold change calculated with respect to control cells. The data presented represent the means of two experiments conducted in triplicate with error bars in the graph indicating the standard error of the mean (SEM). Statistical significance was tested by one-way ANOVA with Dunnett’s correction: *, p≤0.05; **, p≤0.01; ***, p≤0.001. (C) Marmoset kidney cell lines or A549 control cells were treated with conditioned medium from 293T cells expressing human IFNB1 or *Callithrix jacchus* IFNA14. Untreated cells served as control. After 24 h, cell lysates were harvested and analyzed by immunoblot using antibodies against the ISG IFITM3 or beta-actin (as loading control). Results were confirmed in a separate experiment.

To analyze if the marmoset cell lines were responsive to other IFN molecules, we cloned human IFNB1 (hIFNB1) and marmoset IFN α14 (CjIFNa14) and prepared conditioned medium by transfection into 293T cells. Treatment of marmoset cell lines with conditioned medium containing either human IFN β or marmoset IFN α14 clearly induced the ISG IFITM3 as shown in Fig. 3b. This indicates that the cells lines are responsive to IFN and that this response is species specific.

### Immortalized marmoset kidney cells lines are susceptible and permissive for virus infection

To analyze the susceptibility of the cell lines for virus infection, we used MLV-based particles pseudotyped with a large panel of viral glycoproteins of rhabdoviral (VSV-G), filoviral (EBOV-GP, MARV-GP), arenaviral (LCMV-GP, LASV-GP, MACV-GP), bunyaviral (LACV-GPC, RVFV-Gn/Gc, SFTSV-Gn/Gc) and orthomyxoviral (WSN-HA/NA) origin. Most of the pseudoparticles infected all three cell lines with high efficiency (Fig. 4). Notable exceptions were particles bearing influenza A virus WSN-HA/NA, which showed reduced infection especially on CajaK-D4 and CajaK-E5 cells, and particles bearing RVFV-Gn/Gc, which only showed low levels of infection in CajaK-C3 cells but no infection of CajaK-D4 and CajaK-E5 cells. In summary, all immortalized cell lines were susceptible to infection with pseudotypes bearing glycoproteins from a wide range of viruses, including several highly pathogenic ones.

**Figure 4:**
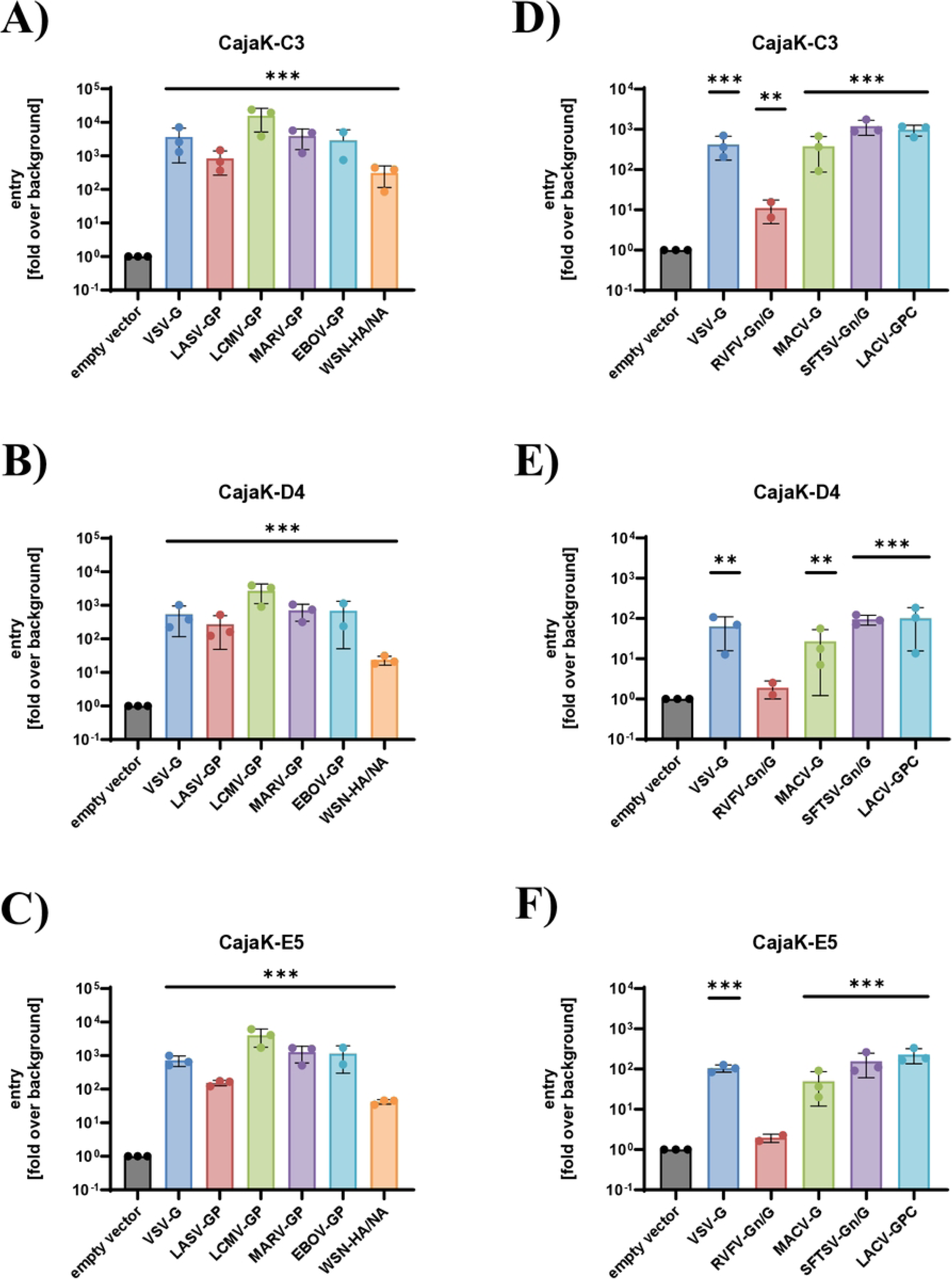
Immortalized marmoset kidney cells are susceptible to entry driven by diverse viral glycoproteins. Marmoset kidney cell lines CajaK-C3 (A, D), CajaK-D4 (B, E) or CajaK-E5 (C, F) were infected with equal volumes of MLV-based particles pseudotyped with glycoproteins of rhabdoviral (VSV-G), filoviral (EBOV-GP, MARV-GP), arenaviral (LCMV-GP, LASV-GP, MACV-GP), bunyaviral (LACV-GPC, RVFV-Gn/Gc, SFTSV-Gn/Gc) and orthomyxoviral (WSN-HA/NA) origin and normalized for comparable luciferase activity upon infection of 293T cells. After 72 h, cell lysates were harvested and luciferase activities measured. The average of the means of two (EBOV-GP, RVFV-Gn/Gc) to three independent experiments (all other) each performed in triplicates is shown. Error bars indicate standard error of the mean (SEM). Statistical significance was tested by one-way ANOVA with Dunnett’s correction: *, p≤0.05; **, p≤0.01; ***, p≤0.001.

Finally, we tested whether the cell lines were permissive to an RNA virus (Zika virus, ZIKV) or a DNA virus (Herpes simplex virus 1, HSV-1). Cells were infected with MOI 1 and virus titer determined in supernatants harvested at regular intervals using focus-formation assay (ZIKV) or plaque assay (HSV-1). Vero76 cells were used as control cells. Both viruses grew on all CajaK cell lines to high levels comparable to Vero76 cells (Fig. 5A, B), indicating efficient and permissive infection. In addition, we used a HSV-1 reporter virus expressing Gaussia luciferase and two closely related reporter viruses of Papiine alphaherpesvirus 2 (PaHV2) and Cercopithecine alphaherpesvirus 2 (CeHV-2) in multicycle infection experiments. Infection war performed at low MOI 0.0004 and replication was quantified using Gaussia luciferase. Here, HSV-1 (Fig. 5C) and PaHV-2 (Fig. 5D) replicated efficiently, while replication of CeHV-2 (Fig. 5E) was clearly reduced. In these experiments, replication was most efficient on CajaK-C3 cells, while Cajak-D4 cells showed reduced replication, which was most pronounced for CeHV-2. In summary, the immortalized CajaK cell lines supported efficient growth of the RNA and DNA viruses tested.

**Figure 5:**
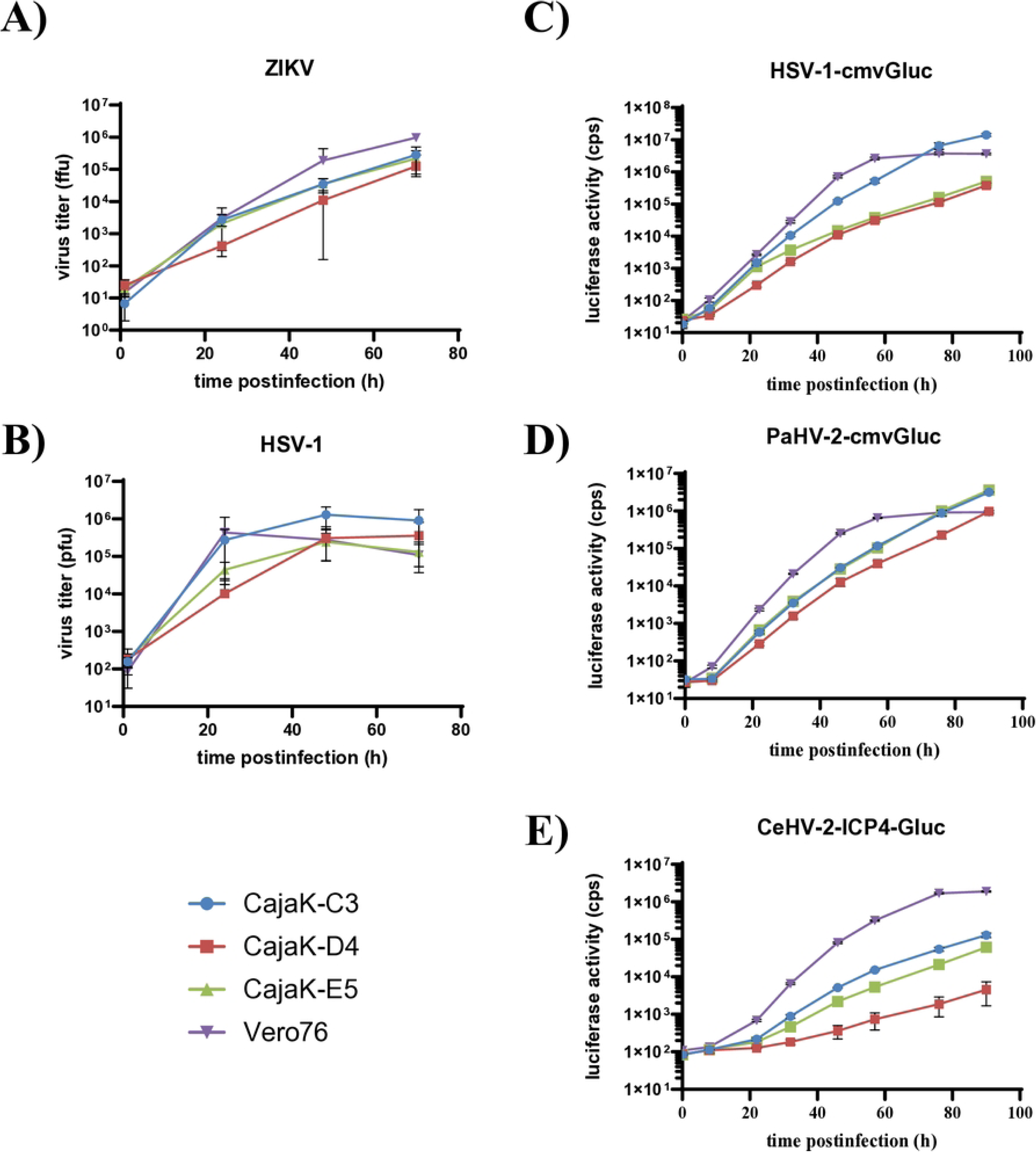
Immortalized marmoset kidney cells support growth of Zika virus and human and primate simplexviruses. (A, B) Infection of cell lines with Zika virus (ZIKV) (A) or Herpes-simplex virus type 1 (HSV-1). Marmoset kidney cell lines of Vero76 control cells were infected with MOI 1 and supernatants harvested at the indicated time points post infection. Virus titers were determined on Vero76 cells using focus-forming assay (ZIKV) or plaque assay (HSV-1). Virus titers are shown as focus forming units (ffu) or plaque forming units (pfu), respectively. The means of two (ZIKV) or three (HSV-1) independent experiment are shown. Error bars indicate standard error of the mean (SEM). (C-E) Multicycle growth curves using reporterviruses HSV-1-cmvGluc, Papiine alphaherpesvirus 2 (PaHV-2) cmvGluc [51] or Cercopithecine alphaherpesvirus 2 (CeHV-2) ICP4-Gluc [52]. Marmoset kidney cell lines or Vero76 control cells were infected at MOI 0.0004 and supernatant was collected at regular intervals. Infection was quantified by measuring Gaussia luciferase activity in the collected supernatants. The results of single representative experiments are shown and were confirmed in two independent experiments. Error bars indicate standard deviation (SD).

## Discussion

In the present work, we describe the generation of three marmoset cell lines of likely pericyte origin, which have a functional IFN system and are susceptible and permissive to infection by multiple viruses. They expand the so far very limited set of immortalized cell lines derived from the common marmoset, a highly valuable NHP model for biomedical research, thus supporting the 3R principles in helping to replace, reduce or refine animal experiments.

So far only few immortalized marmoset cell lines have been described and none was derived from kidney tissue [22]. Our cell lines, derived from marmoset kidney tissue and immortalized by transduction of SV40 large T, therefore markedly expand the repertoire of available marmoset cell lines. Based on RNAseq analysis and comparison with marker genes (COX4I1) and expression profiles derived from human scRNAseq analyses [39, 58], we identified these lines as pericyte-derived, mural cells of the microvasculature involved in maintenance of vessel integrity and regulation of vascular integrity [59]. All three cell lines share similar expression profiles, but clearly have distinct identities as revealed by PCA. These differences may be derived from the phenotypic and transcriptomic diversity of cell types in tissues, but may also be caused in part by expression of the immortalization gene SV40 large T.

The IFN pathway serves as a crucial innate barrier that impacts viral replication. However, many cell lines, for example Vero cells, derived from cancerous tissue lack a functional IFN system [60]. We could show that central steps such as sensing of virus and induction of ISGs are functional in our cell lines. Interestingly, initial treatment with pan-IFN, a chimeric molecule based on human IFNα2 and α4, did not induce the ISG MX1. This may be due to low responsiveness to IFNα2, as has been reported for marmoset hepatocytes treated with human or marmoset IFNα2 [57]. A reason for this may be a general high ISG expression levels in marmoset hepatocytes. Indeed, when analyzed in immunoblot, we observed high levels of IFITM3 expression in our cell lines, as compared to the highly IFN responsive human A549 cell line. However, upon treatment with IFNB1 or cjIFNa14 we clearly noted induction of IFITM3 expression in our marmoset cell lines. In summary, crucial steps of the IFN system are functional in the newly described cell lines.

All cell lines were susceptible to entry driven by glycoproteins of multiple human pathogenic viruses. This includes glycoproteins from viruses causing hemorrhagic fevers, such as LASV, EBOV and MARV, for which marmoset models have already been established [5–7], where cell lines may help to reduce and refine animal experiments. Further, infection by LCMV was found to cause hepatitis in marmosets [61, 62]. The cell lines were also susceptible to entry driven by the glycoproteins of MACV or SFTSV, for which marmoset models have not been described so far. Finally, we could demonstrate that the cell lines were permissive to infection by ZIKV and HSV-1 and related primate simplexviruses. Notably, HSV-1 causes lethal systemic infections in common marmosets after epizootic transmission likely from humans [14, 16, 63] and in a recent outbreak in black-tufted marmosets (*Callithrix penicillata*) virus could be detected in kidney among other organs [64], suggesting that our cell lines model natural target cells.

In summary, we established immortalized kidney cell lines with pericyte identity, which retain a functional IFN system and are permissive to infection with ZIKV, HSV-1 and primate simplexviruses and likely a wide range of other viruses. This broadens the available base of marmoset cell lines and contributes valuable tools for translational research or comparative infection research.

## Acknowledgments

We thank Martina Bleyer (German Primate Center, Göttingen, Germany) for providing the kidney tissue sample, Berit Roshani and Katharina Decker (German Primate Center, Göttingen, Germany) for help with flow cytometry and Gabriela Salinas-Riester and Maren Sitte (Core Unit NGS-Integrative Genomics NIG, Göttingen, Germany) for RNA-seq analysis. We thank Greg Smith (Northwestern University Feinberg School of Medicine, Chicago, USA), Thomas von Hahn (Leibniz University Hannover, Germany), Andrea Maisner (Philipps University of Marburg, Germany), Graham Simmons (UCSF, San Francisco, USA), Birke Bartosch (Lyon, France), Michael Farzan (The Scripps Research Institute, USA), David Brown and Matthew Jones (Public Health England, UK) for their kind gift of material.

## Supporting Information

S1 Fig PCA analysis

S2 Table full gene list normalized counts TPM

S3 Table expression of kidney marker in normalized counts TPM S4 Table expression of pericyte marker in normalized counts TPM S5 Table expression of fibroblast marker in normalized counts TPM S6 Raw images for blots in Fig. 1c and Fic. 3c

## References

1. Herron ICT, Laws TR, Nelson M. Marmosets as models of infectious diseases. Front Cell Infect Microbiol. 2024;14:1340017. Epub 20240223. doi: 10.3389/fcimb.2024.1340017. PubMed PMID: 38465237; PubMed Central PMCID: PMCPMC10921895.

2. Perez-Cruz C, Rodriguez-Callejas JD. The common marmoset as a model of neurodegeneration. Trends Neurosci. 2023;46(5):394–409. Epub 20230310. doi: 10.1016/j.tins.2023.02.002. PubMed PMID: 36907677.

3. Mansfield K. Marmoset models commonly used in biomedical research. Comp Med. 2003;53(4):383–92. PubMed PMID: 14524414.

4. Carrion R, Jr., Patterson JL. An animal model that reflects human disease: the common marmoset (Callithrix jacchus). Curr Opin Virol. 2012;2(3):357–62. Epub 20120314. doi: 10.1016/j.coviro.2012.02.007. PubMed PMID: 22709521; PubMed Central PMCID: PMCPMC3378983.

5. Carrion R, Jr., Brasky K, Mansfield K, Johnson C, Gonzales M, Ticer A, et al. Lassa virus infection in experimentally infected marmosets: liver pathology and immunophenotypic alterations in target tissues. J Virol. 2007;81(12):6482–90. Epub 20070404. doi: 10.1128/JVI.02876-06. PubMed PMID: 17409137; PubMed Central PMCID: PMCPMC1900113.

6. Lukashevich IS, Carrion R, Jr., Salvato MS, Mansfield K, Brasky K, Zapata J, et al. Safety, immunogenicity, and efficacy of the ML29 reassortant vaccine for Lassa fever in small non-human primates. Vaccine. 2008;26(41):5246–54. Epub 20080808. doi: 10.1016/j.vaccine.2008.07.057. PubMed PMID: 18692539; PubMed Central PMCID: PMCPMC2582173.

7. Carrion R, Jr., Ro Y, Hoosien K, Ticer A, Brasky K, de la Garza M, et al. A small nonhuman primate model for filovirus-induced disease. Virology. 2011;420(2):117–24. Epub 20110928. doi: 10.1016/j.virol.2011.08.022. PubMed PMID: 21959017; PubMed Central PMCID: PMCPMC3195836.

8. Chiu CY, Sanchez-San Martin C, Bouquet J, Li T, Yagi S, Tamhankar M, et al. Experimental Zika Virus Inoculation in a New World Monkey Model Reproduces Key Features of the Human Infection. Sci Rep. 2017;7(1):17126. Epub 20171207. doi: 10.1038/s41598-017-17067-w. PubMed PMID: 29215081; PubMed Central PMCID: PMCPMC5719425.

9. Seferovic M, Sanchez-San Martin C, Tardif SD, Rutherford J, Castro ECC, Li T, et al. Experimental Zika Virus Infection in the Pregnant Common Marmoset Induces Spontaneous Fetal Loss and Neurodevelopmental Abnormalities. Sci Rep. 2018;8(1):6851. Epub 20180501. doi: 10.1038/s41598-018-25205-1. PubMed PMID: 29717225; PubMed Central PMCID: PMCPMC5931554.

10. Greenough TC, Carville A, Coderre J, Somasundaran M, Sullivan JL, Luzuriaga K, et al. Pneumonitis and multi-organ system disease in common marmosets (Callithrix jacchus) infected with the severe acute respiratory syndrome-associated coronavirus. Am J Pathol. 2005;167(2):455–63. doi: 10.1016/S0002-9440(10)62989-6. PubMed PMID: 16049331; PubMed Central PMCID: PMCPMC1603565.

11. Singh DK, Singh B, Ganatra SR, Gazi M, Cole J, Thippeshappa R, et al. Responses to acute infection with SARS-CoV-2 in the lungs of rhesus macaques, baboons and marmosets. Nat Microbiol. 2021;6(1):73–86. Epub 20201218. doi: 10.1038/s41564-020-00841-4. PubMed PMID: 33340034; PubMed Central PMCID: PMCPMC7890948.

12. Falzarano D, de Wit E, Feldmann F, Rasmussen AL, Okumura A, Peng X, et al. Infection with MERS-CoV causes lethal pneumonia in the common marmoset. PLoS Pathog. 2014;10(8):e1004250. Epub 20140821. doi: 10.1371/journal.ppat.1004250. PubMed PMID: 25144235; PubMed Central PMCID: PMCPMC4140844.

13. tHart BA, Kap YS, Morandi E, Laman JD, Gran B. EBV Infection and Multiple Sclerosis: Lessons from a Marmoset Model. Trends Mol Med. 2016;22(12):1012–24. Epub 20161108. doi: 10.1016/j.molmed.2016.10.007. PubMed PMID: 27836419.

14. Matz-Rensing K, Jentsch KD, Rensing S, Langenhuyzen S, Verschoor E, Niphuis H, et al. Fatal Herpes simplex infection in a group of common marmosets (Callithrix jacchus). Vet Pathol. 2003;40(4):405–11. doi: 10.1354/vp.40-4-405. PubMed PMID: 12824512.

15. Hatt JM, Grest P, Posthaus H, Bossart W. Serologic survey in a colony of captive common marmosets (Callithrix jacchus) after infection with herpes simplex type 1-like virus. J Zoo Wildl Med. 2004;35(3):387–90. doi: 10.1638/03-041. PubMed PMID: 15526895.

16. Imura K, Chambers JK, Uchida K, Nomura S, Suzuki S, Nakayama H, et al. Herpes simplex virus type 1 infection in two pet marmosets in Japan. J Vet Med Sci. 2014;76(12):1667–70. Epub 20140911. doi: 10.1292/jvms.14-0374. PubMed PMID: 25649955; PubMed Central PMCID: PMCPMC4300388.

17. Russell WMS, Burch RL. The principles of humane experimental technique. London: Methuen; 1959.

18. Petkov S, Kahland T, Shomroni O, Lingner T, Salinas G, Fuchs S, et al. Immortalization of common marmoset monkey fibroblasts by piggyBac transposition of hTERT. PLoS One. 2018;13(9):e0204580. Epub 2018/09/28. doi: 10.1371/journal.pone.0204580. PubMed PMID: 30261016; PubMed Central PMCID: PMCPMC6160115.

19. Orimoto A, Shinohara H, Eitsuka T, Nakagawa K, Sasaki E, Kiyono T, et al. Immortalization of common marmoset-derived fibroblasts via expression of cell cycle regulators using the piggyBac transposon. Tissue Cell. 2022;77:101848. Epub 20220602. doi: 10.1016/j.tice.2022.101848. PubMed PMID: 35714414.

20. Husen B, Lieder K, Marten A, Jurdzinski A, Fuhrmann K, Petry H, et al. Immortalisation of ovarian granulosa and Theca cells of the marmoset monkey Calllithrix jacchus. ALTEX. 2002;19 Suppl 1:64–72. PubMed PMID: 12096332.

21. Guo Z, Jing R, Rao Q, Zhang L, Gao Y, Liu F, et al. Immortalized common marmoset (Callithrix jacchus) hepatic progenitor cells possess bipotentiality in vitro and in vivo. Cell Discov. 2018;4:23. Epub 20180515. doi: 10.1038/s41421-018-0020-7. PubMed PMID: 29796307; PubMed Central PMCID: PMCPMC5951880.

22. Bayurova E, Zhitkevich A, Avdoshina D, Kupriyanova N, Kolyako Y, Kostyushev D, et al. Common Marmoset Cell Lines and Their Applications in Biomedical Research. Cells. 2023;12(16). Epub 20230808. doi: 10.3390/cells12162020. PubMed PMID: 37626830; PubMed Central PMCID: PMCPMC10453182.

23. Bartosch B, Dubuisson J, Cosset FL. Infectious hepatitis C virus pseudo-particles containing functional E1-E2 envelope protein complexes. J Exp Med. 2003;197(5):633–42. doi: 10.1084/jem.20021756. PubMed PMID: 12615904; PubMed Central PMCID: PMCPMC2193821.

24. Reiter S, Gartner S, Decker K, Pohlmann S, Winkler M. Development of immortalized rhesus macaque kidney cells supporting infection with a panel of viruses. PLoS One. 2023;18(5):e0284048. Epub 2023/05/05. doi: 10.1371/journal.pone.0284048. PubMed PMID: 37146034; PubMed Central PMCID: PMCPMC10162512.

25. Reiter S, Sun T, Gartner S, Pohlmann S, Winkler M. Development of rhesus macaque astrocyte cell lines supporting infection with a panel of viruses. PLoS One. 2024;19(5):e0303059. Epub 20240514. doi: 10.1371/journal.pone.0303059. PubMed PMID: 38743751; PubMed Central PMCID: PMCPMC11093292.

26. Marzi A, Gramberg T, Simmons G, Moller P, Rennekamp AJ, Krumbiegel M, et al. DC-SIGN and DC-SIGNR interact with the glycoprotein of Marburg virus and the S protein of severe acute respiratory syndrome coronavirus. J Virol. 2004;78(21):12090–5. doi: 10.1128/JVI.78.21.12090-12095.2004. PubMed PMID: 15479853; PubMed Central PMCID: PMCPMC523257.

27. Fouchier RA, Meyer BE, Simon JH, Fischer U, Malim MH. HIV-1 infection of non-dividing cells: evidence that the amino-terminal basic region of the viral matrix protein is important for Gag processing but not for post-entry nuclear import. EMBO J. 1997;16(15):4531–9. doi: 10.1093/emboj/16.15.4531. PubMed PMID: 9303297; PubMed Central PMCID: PMCPMC1170079.

28. Chaipan C, Kobasa D, Bertram S, Glowacka I, Steffen I, Tsegaye TS, et al. Proteolytic activation of the 1918 influenza virus hemagglutinin. J Virol. 2009;83(7):3200–11. doi: 10.1128/JVI.02205-08. PubMed PMID: 19158246; PubMed Central PMCID: PMCPMC2655587.

29. Simmons G, Reeves JD, Grogan CC, Vandenberghe LH, Baribaud F, Whitbeck JC, et al. DC-SIGN and DC-SIGNR bind ebola glycoproteins and enhance infection of macrophages and endothelial cells. Virology. 2003;305(1):115–23. doi: 10.1006/viro.2002.1730. PubMed PMID: 12504546.

30. Radoshitzky SR, Abraham J, Spiropoulou CF, Kuhn JH, Nguyen D, Li W, et al. Transferrin receptor 1 is a cellular receptor for New World haemorrhagic fever arenaviruses. Nature. 2007;446(7131):92–6. doi: 10.1038/nature05539. PubMed PMID: 17287727; PubMed Central PMCID: PMCPMC3197705.

31. Plassmeyer ML, Soldan SS, Stachelek KM, Martin-Garcia J, Gonzalez-Scarano F. California serogroup Gc (G1) glycoprotein is the principal determinant of pH-dependent cell fusion and entry. Virology. 2005;338(1):121–32. doi: 10.1016/j.virol.2005.04.026. PubMed PMID: 15923017.

32. Habjan M, Penski N, Spiegel M, Weber F. T7 RNA polymerase-dependent and - independent systems for cDNA-based rescue of Rift Valley fever virus. J Gen Virol. 2008;89(Pt 9):2157–66. doi: 10.1099/vir.0.2008/002097-0. PubMed PMID: 18753225.

33. Hofmann H, Li X, Zhang X, Liu W, Kuhl A, Kaup F, et al. Severe fever with thrombocytopenia virus glycoproteins are targeted by neutralizing antibodies and can use DC-SIGN as a receptor for pH-dependent entry into human and animal cell lines. J Virol. 2013;87(8):4384–94. Epub 20130206. doi: 10.1128/JVI.02628-12. PubMed PMID: 23388721; PubMed Central PMCID: PMCPMC3624395.

34. Niwa H, Yamamura K, Miyazaki J. Efficient selection for high-expression transfectants with a novel eukaryotic vector. Gene. 1991;108(2):193–9. doi: 10.1016/0378-1119(91)90434-d. PubMed PMID: 1660837.

35. Dirks WG, Drexler HG. STR DNA typing of human cell lines: detection of intra-and interspecies cross-contamination. Methods Mol Biol. 2013;946:27–38. doi: 10.1007/978-1-62703-128-8_3. PubMed PMID: 23179824.

36. Schindelin J, Arganda-Carreras I, Frise E, Kaynig V, Longair M, Pietzsch T, et al. Fiji: an open-source platform for biological-image analysis. Nat Methods. 2012;9(7):676–82. doi: 10.1038/nmeth.2019. PubMed PMID: 22743772.

37. Kolde R. pheatmap: pretty heatmaps 2019. Available from: https://cran.r-project.org/web/packages/pheatmap/pheatmap.pdf.

38. Hu K. Become Competent in Generating RNA-Seq Heat Maps in One Day for Novices Without Prior R Experience. Methods Mol Biol. 2021;2239:269–303. doi: 10.1007/978-1-0716-1084-8_17. PubMed PMID: 33226625.

39. Subramanian A, Sidhom EH, Emani M, Vernon K, Sahakian N, Zhou Y, et al. Single cell census of human kidney organoids shows reproducibility and diminished off-target cells after transplantation. Nat Commun. 2019;10(1):5462. Epub 20191129. doi: 10.1038/s41467-019-13382-0. PubMed PMID: 31784515; PubMed Central PMCID: PMCPMC6884507.

40. Ma Y, Eizenberg-Magar I, Antebi Y. EasyFlow: An open-source, user-friendly cytometry analyzer with graphic user interface (GUI). PLoS One. 2024;19(11):e0308873. Epub 20241113. doi: 10.1371/journal.pone.0308873. PubMed PMID: 39536028; PubMed Central PMCID: PMCPMC11560029.

41. Holzinger D, Jorns C, Stertz S, Boisson-Dupuis S, Thimme R, Weidmann M, et al. Induction of MxA gene expression by influenza A virus requires type I or type III interferon signaling. J Virol. 2007;81(14):7776–85. doi: 10.1128/JVI.00546-06. PubMed PMID: 17494065; PubMed Central PMCID: PMCPMC1933351.

42. Hayashi F, Means TK, Luster AD. Toll-like receptors stimulate human neutrophil function. Blood. 2003;102(7):2660–9. doi: 10.1182/blood-2003-04-1078. PubMed PMID: 12829592.

43. Manthey HD, Thomas AC, Shiels IA, Zernecke A, Woodruff TM, Rolfe B, et al. Complement C5a inhibition reduces atherosclerosis in ApoE-/- mice. FASEB J. 2011;25(7):2447–55. doi: 10.1096/fj.10-174284. PubMed PMID: 21490292.

44. Livak KJ, Schmittgen TD. Analysis of relative gene expression data using real-time quantitative PCR and the 2(-Delta Delta C(T)) Method. Methods. 2001;25(4):402–8. doi: 10.1006/meth.2001.1262. PubMed PMID: 11846609.

45. Longo PA, Kavran JM, Kim MS, Leahy DJ. Transient mammalian cell transfection with polyethylenimine (PEI). Methods Enzymol. 2013;529:227–40. doi: 10.1016/B978-0-12-418687-3.00018-5. PubMed PMID: 24011049; PubMed Central PMCID: PMCPMC4012321.

46. Berger Rentsch M, Zimmer G. A vesicular stomatitis virus replicon-based bioassay for the rapid and sensitive determination of multi-species type I interferon. PLoS One. 2011;6(10):e25858. Epub 20111005. doi: 10.1371/journal.pone.0025858. PubMed PMID: 21998709; PubMed Central PMCID: PMCPMC3187809.

47. Eckert N, Wrensch F, Gartner S, Palanisamy N, Goedecke U, Jager N, et al. Influenza A virus encoding secreted Gaussia luciferase as useful tool to analyze viral replication and its inhibition by antiviral compounds and cellular proteins. PLoS One. 2014;9(5):e97695. doi: 10.1371/journal.pone.0097695. PubMed PMID: 24842154; PubMed Central PMCID: PMCPMC4026478.

48. Wrensch F, Winkler M, Pohlmann S. IFITM proteins inhibit entry driven by the MERS-coronavirus spike protein: evidence for cholesterol-independent mechanisms. Viruses. 2014;6(9):3683–98. doi: 10.3390/v6093683. PubMed PMID: 25256397; PubMed Central PMCID: PMCPMC4189045.

49. Braun E, Hotter D, Koepke L, Zech F, Gross R, Sparrer KMJ, et al. Guanylate-Binding Proteins 2 and 5 Exert Broad Antiviral Activity by Inhibiting Furin-Mediated Processing of Viral Envelope Proteins. Cell Rep. 2019;27(7):2092–104 e10. doi: 10.1016/j.celrep.2019.04.063. PubMed PMID: 31091448.

50. Richards AL, Sollars PJ, Smith GA. New tools to convert bacterial artificial chromosomes to a self-excising design and their application to a herpes simplex virus type 1 infectious clone. BMC Biotechnol. 2016;16(1):64. Epub 2016/09/02. doi: 10.1186/s12896-016-0295-4. PubMed PMID: 27580861; PubMed Central PMCID: PMCPMC5006514.

51. Rahman Siregar A, Gartner S, Gotting J, Stegen P, Kaul A, Schulz TF, et al. A Recombinant System and Reporter Viruses for Papiine Alphaherpesvirus 2. Viruses. 2022;14(1). Epub 2022/01/23. doi: 10.3390/v14010091. PubMed PMID: 35062295; PubMed Central PMCID: PMCPMC8778148.

52. Chukhno E, Gartner S, Rahman Siregar A, Mehr A, Wende M, Petkov S, et al. A Fosmid-Based System for the Generation of Recombinant Cercopithecine Alphaherpesvirus 2 Encoding Reporter Genes. Viruses. 2019;11(11). doi: 10.3390/v11111026. PubMed PMID: 31694178; PubMed Central PMCID: PMCPMC6893520.

53. Tannous BA. Gaussia luciferase reporter assay for monitoring biological processes in culture and in vivo. Nat Protoc. 2009;4(4):582–91. doi: 10.1038/nprot.2009.28. PubMed PMID: 19373229; PubMed Central PMCID: PMCPMC2692611.

54. Motulsky HJ, Head T, Clarke PBS. Analyzing lognormal data: A nonmathematical practical guide. Pharmacol Rev. 2025;77(3):100049. Epub 20250225. doi: 10.1016/j.pharmr.2025.100049. PubMed PMID: 40153903; PubMed Central PMCID: PMCPMC12163497.

55. Ronni T, Matikainen S, Sareneva T, Melen K, Pirhonen J, Keskinen P, et al. Regulation of IFN-alpha/beta, MxA, 2’,5’-oligoadenylate synthetase, and HLA gene expression in influenza A-infected human lung epithelial cells. J Immunol. 1997;158(5):2363–74. PubMed PMID: 9036986.

56. Hoffmann M, Wu YJ, Gerber M, Berger-Rentsch M, Heimrich B, Schwemmle M, et al. Fusion-active glycoprotein G mediates the cytotoxicity of vesicular stomatitis virus M mutants lacking host shut-off activity. J Gen Virol. 2010;91(Pt 11):2782–93. doi: 10.1099/vir.0.023978-0. PubMed PMID: 20631091.

57. Chavez D, Guerra B, Lanford RE. Antiviral activity and host gene induction by tamarin and marmoset interferon-alpha and interferon-gamma in the GBV-B primary hepatocyte culture model. Virology. 2009;390(2):186–96. Epub 2009/06/09. doi: 10.1016/j.virol.2009.05.005. PubMed PMID: 19501869; PubMed Central PMCID: PMCPMC2753388.

58. Balzer MS, Rohacs T, Susztak K. How Many Cell Types Are in the Kidney and What Do They Do? Annu Rev Physiol. 2022;84:507–31. Epub 20211129. doi: 10.1146/annurev-physiol-052521-121841. PubMed PMID: 34843404; PubMed Central PMCID: PMCPMC9233501.

59. Tuleta I, Frangogiannis NG. Pericytes in tissue fibrosis. Am J Physiol Cell Physiol. 2025;329(3):C868-C86. Epub 20250812. doi: 10.1152/ajpcell.00403.2025. PubMed PMID: 40796212; PubMed Central PMCID: PMCPMC12439126.

60. Osada N, Kohara A, Yamaji T, Hirayama N, Kasai F, Sekizuka T, et al. The genome landscape of the african green monkey kidney-derived vero cell line. DNA Res. 2014;21(6):673–83. doi: 10.1093/dnares/dsu029. PubMed PMID: 25267831; PubMed Central PMCID: PMCPMC4263300.

61. Stephensen CB, Jacob JR, Montali RJ, Holmes KV, Muchmore E, Compans RW, et al. Isolation of an arenavirus from a marmoset with callitrichid hepatitis and its serologic association with disease. J Virol. 1991;65(8):3995–4000. doi: 10.1128/JVI.65.8.3995-4000.1991. PubMed PMID: 1712856; PubMed Central PMCID: PMCPMC248829.

62. Asper M, Hofmann P, Osmann C, Funk J, Metzger C, Bruns M, et al. First outbreak of callitrichid hepatitis in Germany: genetic characterization of the causative lymphocytic choriomeningitis virus strains. Virology. 2001;284(2):203–13. doi: 10.1006/viro.2001.0909. PubMed PMID: 11384220.

63. Juan-Salles C, Ramos-Vara JA, Prats N, Sole-Nicolas J, Segales J, Marco AJ. Spontaneous herpes simplex virus infection in common marmosets (Callithrix jacchus). J Vet Diagn Invest. 1997;9(3):341–5. doi: 10.1177/104063879700900325. PubMed PMID: 9249183.

64. Garcia-Oliveira GF, Frasson Biccas M, Jacob D, Oliveira MA, Paschoal AMO, Alves PA, et al. Human Herpesvirus 1 Associated with Epizootics in Belo Horizonte, Minas Gerais, Brazil. Viruses. 2025;17(5). Epub 20250430. doi: 10.3390/v17050660. PubMed PMID: 40431671; PubMed Central PMCID: PMCPMC12115514.

